# *Lhr* and *Hmr* are required for sister chromatid detachment during anaphase but not for centromere function

**DOI:** 10.1101/178046

**Authors:** Jacob A. Blum, Silvia Bonaccorsi, Marta Marzullo, Valeria Palumbo, Yukiko M. Yamashita, Daniel A. Barbash, Maurizio Gatti

**Author notes:** Correspondence: Daniel A. Barbash < > Maurizio Gatti < > Dipartimento di Biologia e Biotecnologie “C. Darwin”, Sapienza, Università di Roma 00185 Roma, Italy. These authors contributed equally (Jacob A. Blum, Silvia Bonaccorsi).

## Abstract

Crosses between *Drosophila melanogaster* females and *Drosophila simulans* males produce hybrid sons that die at the larval stage. This hybrid lethality is suppressed by loss-of-function mutations in the *D. melanogaster Hybrid male rescue* (*Hmr*) or in the *D. simulans Lethal hybrid rescue* (*Lhr)* genes. Previous studies have shown that Hmr and Lhr interact with heterochromatin proteins and suppress expression of transposable elements within *D. melanogaster*. It also has been proposed that Hmr and Lhr function at the centromere. We examined mitotic divisions in larval brains from *Hmr* and *Lhr* single mutants and *Hmr; Lhr* double mutants in *D. melanogaster*. In none of the mutants did we observe defects in metaphase chromosome alignment or hyperploid cells, which are hallmarks of centromere or kinetochore dysfunction. In addition, we found that Hmr-HA and Lhr-HA do not localize to centromeres either during interphase or mitotic division. However, all mutants displayed anaphase bridges and chromosome aberrations resulting from the breakage of these bridges, predominantly at the euchromatin-heterochromatin junction. The few dividing cells present in hybrid males showed irregularly condensed chromosomes with fuzzy and often closely apposed sister chromatids. Despite this defect in condensation, chromosomes in hybrids managed to align on the metaphase plate and undergo anaphase. We conclude that there is no evidence for a centromeric function of Hmr and Lhr within *D. melanogaster* nor for a centromere defect causing hybrid lethality. Instead we find that *Hmr* and *Lhr* are required in *D. melanogaster* for detachment of sister chromatids during anaphase.

## Introduction

The reduced fertility and viability of interspecific hybrids are widely observed causes of reproductive isolation that contribute to speciation. According to the classical Dobzhansky-Muller (D-M) model of hybrid incompatibility (HI), two or more loci that had independently diverged in nascent species can lead to deleterious HI effects when combined in interspecific hybrids. Although the D-M model is generally accepted and numerous HI genes have been identified, the cytological and molecular mechanisms underlying HI are still poorly understood (reviewed in Presgraves, 2010; Maheshwari and Barbash, 2011).

Hybrids produced with *Drosophila melanogaster* offer strong opportunities to investigate mechanisms that cause HI (Barbash 2010). Crosses between *D. melanogaster* females and *Drosophila simulans* males produce viable but sterile females and no males, which die at the larval stage. This hybrid lethality is suppressed by loss of function mutations in the *D. melanogaster Hybrid male rescue* (*Hmr*) gene or in the *D. simulans Lethal hybrid rescue* (*Lhr)* gene (Maheshwari and Barbash, 2011). The hybrid phenotype of these mutations indicates that hybrid lethality is caused by the wild type alleles of these genes, which are therefore functioning as gain-of-function mutations in the hybrid background. Hybrid male larvae from crosses between *D. melanogaster* females and *D. simulans* males die prior to pupal differentiation and exhibit small brains and an almost complete absence of imaginal discs. Most cells in the brains of these larvae appear to be arrested either in G1 or G2, and the few dividing cells display defects in chromosome morphology (Orr et al., 1997; Bolkan et al., 2007).

A crucial step to elucidate how the interaction of *Hmr* and *Lhr* leads to hybrid lethality is to understand first their biological roles in each of the two species. Studies carried out in *D. melanogaster* have shown that neither *Hmr* nor the orthologue of *D. simulans Lhr* (henceforth designated *Lhr* without specifying that it is the *D. melanogaster* gene) is an essential gene. Flies homozygous for null mutations in either *Hmr* or *Lhr* are viable but have reduced female fertility (Aruna et al., 2009; Satyaki et al., 2014).

The Hmr and Lhr proteins are chromosome-associated and enriched in the heterochromatin. In interphase embryonic cells, both proteins largely co-localize with the heterochromatin markers HP1a and H3K9me2 (histone H3 di-methylated at K9) (Maheshwari and Barbash, 2012; Satyaki et al., 2014). In polytene chromosomes, Hmr and Lhr are enriched in both the alpha- and beta-heterochromatin of the chromocenter, in a few euchromatic bands, and at the telomeres (Thomae et al., 2013; Satyaki et al., 2014). Alpha heterochromatin occupies a small area in the middle of the chromocenter and contains mitotic heterochromatin and satellite DNAs, which are severely under-replicated in polytene chromosomes. Alpha heterochromatin is connected to the euchromatic chromosome arms by *beta* heterochromain, which is enriched in diverse arrays of unique and repetitive DNA sequences but not in satellite DNA (Miklos and Cotsel, 1990; Gatti and Pimpinelli, 1992). Consistent with their heterochromatic and telomeric localizations, Hmr and Lhr associate with heterochromatin protein 1a (HP1a), and Hmr and Lhr interact with each other in the yeast two-hybrid assay, suggesting that the three proteins are part of a complex within which Hmr and Lhr interact directly (Thomae et al., 2013; Alekseyenko et al., 2014; Satyaki et al., 2014).

Thomae et al. (2013) proposed that Hmr and Lhr are centromere proteins. This suggestion was based on three main findings. They reported that in interphase imaginal disc cells, Hmr and Lhr co-localize with heterochromatic regions that are largely coincident with those immunostained for the centromere markers Cid and Cenp-C (Thomae et al., 2013). Using tandem co-purification experiments followed by mass spectrometry, and additional co-precipitation experiments, they identified 60 Hmr-Lhr interacting proteins including Cenp-C that is a centromere-specific component (Heeger et al., 2005), as well as HP1 and HP6/Umbrea that are enriched in centromeric heterochromatin (Greil et al., 2007; Ross at al., 2013). They also observed lagging chromosomes in anaphases of Hmr- and Lhr-depleted cells (Thomae et al., 2013). Several aspects of their report, however, leave open alternative interpretations about Hmr and Lhr function. First, many of the co-purifying proteins they identified have non-centromeric functions. For example, HP1a and HP6/Umbrea localize also in non-centromeric heterochromatic regions and at telomeres, and HP1a has been shown to prevent telomere fusion in somatic cells (Fanti et al, 1998; Joppich et al., 2009; Vermaak and Malik, 2009; Elgin and Reuter, 2013). In addition, three identified proteins are components of the *Drosophila* telomere-capping complex (Ver, Moi and CG30007/Tea), identified by lethal mutations that cause frequent telomeric fusions in larval brain cells (Raffa et al., 2009; Raffa et al 2010; Cicconi et al., 2016; Zhang et al., 2016). Thus, the interactions between Hmr-Lhr and proteins such as HP1a, HP6/Umbrea do not necessarily occur at centromeres. Second, centromeric localization of Hmr and Lhr was not observed in metaphase chromosomes (Thomae et al., 2013), nor did Lhr co-localize with Cid in embryonic interphase nuclei (Maheshwari et al, 2012). Third, the centromeric role of Hmr and Lhr proposed by Thomae et al is unclear, because they found that loss of neither Hmr nor Lhr affects centromeric localization of essential centromere/kinetochore components including Cid, Cenp-C, Ndc80, Incenp, Polo and Rod. We therefore investigated here using extensive cytological analysis of larval brain cells whether *Hmr* and *Lhr* affect centromere function, or potentially a different aspect of chromosome segregation. We found that these mutants exhibit very low levels of telomeric fusions. However, they displayed relatively high frequencies of incomplete chromosome breaks, namely broken chromosomes without the corresponding fragment or complete chromosome complements plus an extra acentric fragment. These two types of chromosome aberrations are likely generated during anaphase (Mengoli et al., 2014). Notably, we did not observe aneuploid cells with unbroken chromosomes in either *Hmr* or *Lhr* mutant brains or failure of the centromeres to separate and move towards the poles. In addition, immunolocalization experiment in larval brain cells showed that neither Hmr-HA nor Lhr-HA localizes to the centromeres throughout the cell cycle. Thus, our results strongly suggest that the Hmr-Lhr complex is not required for centromere or kinetochore function.

## Materials and Methods

### Drosophila strains and crosses

*Hmr*^*3*^, also called *Hmr*^*EY12237*^, carries a {EPgy2} insertion within the gene and is a null allele by genetic criteria (Aruna et al., 2009). *Lhr*^*KO*^, generated by a targeted *w*^*+*^ insertion within the gene, carries a 26 bp deletion of the coding sequence and is also a null mutation (Satyaki et al., 2014). To generate the double mutant we crossed *Hmr*^*3*^*/Y; Lhr*^*KO*^*/CyO-Tb* males to *Hmr*^*3*^*/FM7-Tb; Lhr*^*KO*^*/CyO-Tb* females; the non-Tb progeny (*Hmr*^*3*^*/Y; Lhr*^*KO*^*/Lhr*^*KO*^ males and *Hmr*^*3*^*/Hmr*^*3*^*; Lhr*^*KO*^*/Lhr*^*KO*^ females) from these crosses were then examined for chromosome integrity and mitotic cell morphology. The *FM7-Tb* and *CyO-Tb* balancers, which carry the dominant larval marker *Tb*, are described in Lattao et al. (2011).

### Chromosome cytology

To analyze chromosome aberrations in metaphases, brains from third-instar larvae were dissected in saline (NaCl 0.7%) and incubated for 1h in saline with colchicine (10^-5^ M); brains were then treated for 8 min with hypotonic solution (0.5% Na Citrate), and squashed in 45% acetic acid under a 20 x 20 mm coverslip. To analyze anaphases and assess mitotic parameters, larval brains were dissected in saline and directly squashed in 45% acetic acid without colchicine and hypotonic pretreatment. Both types of chromosome squashes were frozen in liquid nitrogen; after flipping off the coverslip slides were air dried and then mounted in Vectashield H-1200 (Vector Laboratories, Burlingame, CA) with DAPI (4,6 diamidino-2-phenylindole) to stain the chromosomes and reduce fluorescence fading.

### Subcellular localization of Hmr-HA and Lhr-HA

The Hmr-HA and Lhr-HA transgenes were described previously (Maheshwari and Barbash 2012; Satyaki et al. 2014). To examine their localization in larval brain cells, brains from crawling third instar larvae were dissected in PBS, transferred to 0.5% Na citrate for 10-20 minutes, and transferred onto a 25μl drop of fixative (4% formaldehyde in PBST). While being fixed (for 4 minutes), the brains were manually dissected into smaller pieces to ensure better spread of cells. After fixation, the tissues were squashed and frozen in liquid nitrogen. After flipping off the coverslips, slides were washed in PBS for ∼1hr, and incubated overnight with primary antibodies (chicken anti-Cid; 1:500, generated at Covance against the peptide AKRAPRPSANNSKSPNDD; and rat anti-HA, 1:100, Sigma-Aldrich) in 3% BSA in PBST at 4°C. The slides were washed in PBS for ∼1 hour and then incubated overnight with secondary antibodies (anti-Chicken AlexaFluor-488, anti-rat AlexaFluor-568) at 4°C. The slides were washed for ∼1hr and mounted in Vectashield H-1200 (Vector Laboratories, Burlingame, CA) with DAPI (4,6 diamidino-2-phenylindole).

### Mitotic chromosome and spindle immunostaining

For immunostaining with anti-tubulin and anti phospho-histone H3 (PH3) antibodies, brains from third instar larvae were dissected in saline, fixed in formaldehyde and squashed as described in Bonaccorsi et al. (2000). For PH3 immunostaining, preparations were incubated overnight at 4°C with a rabbit anti-PH3 (Ser10) antibody (Upstate Biotechnology, Lake Placid, NY) diluted 1:100 in PBS with 5% goat serum. The anti-PH3 antibody was detected by a 1-hour incubation at room temperature with an Alexa 555-conjugated anti-rabbit IgG (Molecular Probes, Eugene, OR) diluted 1:300 in PBS. For tubulin immunostaining slides were incubated overnight at 4°C with an anti-α tubulin monoclonal (DM1A diluted 1:100; Sigma), which was detected by a 1-hour incubation at room temperature with FITC-conjugated anti-mouse (1:100, Jackson Laboratories, Bar Harbor, ME) diluted in PBS. All cytological preparations were mounted in Vectashield H-1200 with DAPI and images were captured with a CoolSnap HQ CCD camera (Photometrics; Tucson, AZ) connected to a Zeiss Axioplan fluorescence microscope equipped with an HBO 100 W mercury lamp.

## Results

### Structure of chromosomes in Hmr and Lhr mutant stocks

Because both *Hmr* and *Lhr* have been implicated in the maintenance of heterochromatin, we first examined the mitotic chromosomes of each mutant for the structure of the heterochromatic regions. This is an important control to exclude the possibility that the chromosome aberration phenotypes that we describe below are due to chromosomal abnormalities that pre-exist in the mutant stocks that we are using. We crossed mutant males with wild type Oregon-R females and then examined the heterozygous progeny for the DAPI banding pattern of larval brain heterochromatin. In late prophase cells of these larval brains the Oregon-R chromosome is paired with its mutant homologue, facilitating a comparison between the heterochromatic regions. Because the Hoechst banding pattern of the Oregon-R heterochromatin (which is identical to that obtained with DAPI) has been carefully characterized (reviewed by Gatti and Pimpinelli, 1992), this analysis permitted us to assess precisely whether the heterochromatic regions of the chromosomes from the *Hmr*^*3*^ and *Lhr*^*KO*^ stocks are different from those of Oregon-R wild type flies. We found that the heterochromatic regions of both mutants are virtually identical to those of Oregon-R, with a minor difference in the most distal fluorescent bands of the 3R heterochromatin in *Lhr*^*KO*^ flies (Figure 1). Although the precise nature of this difference is not clear, it is certainly insufficient to account for the range of chromosome aberrations described below (Figure 2, tables 1 and 2).

**Figure 1.**
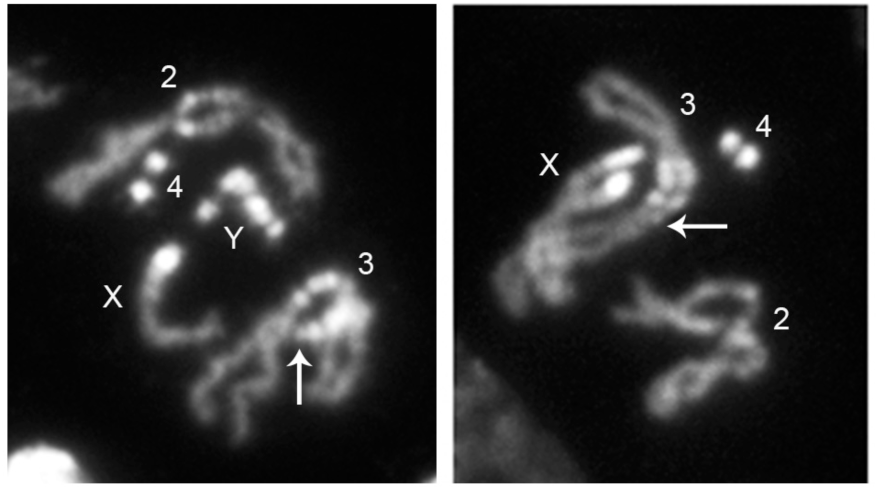
*Lhr*^*KO*^ mutants exhibit a minor difference in the most distal fluorescent bands of the 3R heterochromatin (arrows) compared to wild type Oregon R flies. The cells shown are male (left) and female (right) late prophases from F1 third instar larvae generated by crosses between Oregon R females and *Lhr*^*KO*^ mutant males.

**Figure 2.**
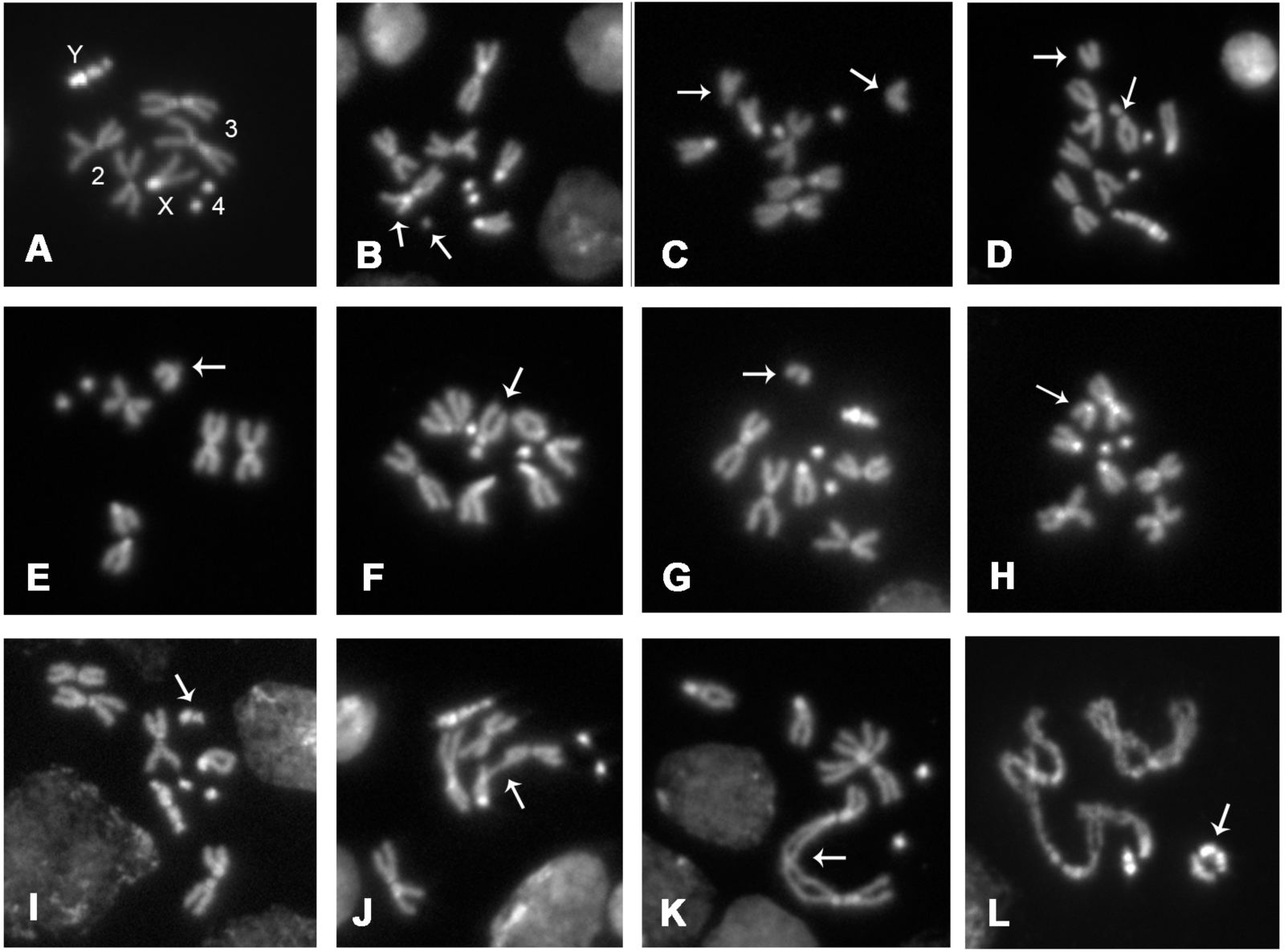
Examples of chromosome aberrations observed in colchicine-treated metaphases from *Lhr* and *Hmr* mutants. A. Male control metaphase; the third chromosomes are easily distinguished from the second chromosomes by the higher fluorescence of their centric heterochromatin. B. Chromatid deletion (arrows). C. Second chromosome ISOB in the centric heterochromatin, probably at the euchromatin-heterochromatin (eu-het) junction (arrows). D. Third chromosome ISOB at the eu-het junction (arrows). E. Incomplete ISOB; a second chromosome (arrow) is broken within the heterochromatin or at the eu-het junction but lacks the corresponding acentric fragment. F. Incomplete ISOB; a third chromosome broken at the eu-het junction (arrow) lacking the corresponding acentric fragment. G, H and I. Metaphases with complete chromosome complements and an additional euchromatic fragment (G; arrow), an additional autosomal arm broken in the heterochromatin (H, arrow), or an additional Y fragment (I, arrow). J. Telomeric fusion involving single chromatids of the X and the second chromosome (arrow). K. Double telomeric fusion involving both sister chromatids of the two third chromosomes (arrow). L. Telomeric fusion leading to a ring Y chromosome (arrow).

**Table 1.** Mutations in *Lhr* and *Hmr* cause chromosome aberrations (CABs)

| Mutant, sex | # of brains | #of cells | CD | ISOBs with F |  |  |  | ISOBs with no F |  |  |  | Extra F |  |  | E | TFs | H | CABs % | TF % |
| --- | --- | --- | --- | --- | --- | --- | --- | --- | --- | --- | --- | --- | --- | --- | --- | --- | --- | --- | --- |
|  |  |  |  | eu | Ah | X | Y | eu | Ah | X | Y | eu | Ah | Y |  |  |  |  |  |
| *Hmr*<sup>3</sup>, f | 14 | 914 | 2 | 6 | 7 | 2 | - | 1 | 13 | 4 | - | 6 | 8 | - | 2 | 8 | 2 | 5.6 | 0.9 |
| *Hmr*<sup>3</sup>, m | 12 | 920 | 1 | 9 | 7 | 4 | 2 | 1 | 15 | 1 | 3 | 2 | 4 | 3 | 1 | 6 | 3 | 5.8 | 0.7 |
| *Hmr*<sup>3/+</sup>, f | 9 | 929 | 0 | 1 | 0 | 1 | - | 0 | 0 | 0 | - | 1 | 1 | - | 0 | 0 | 0 | 0.5 | 0 |
| *Lhr*<sup>KO</sup>, f | 18 | 1,404 | 6 | 4 | 14 | 5 | - | 2 | 19 | 6 | - | 7 | 10 | - | 0 | 6 | 2 | 5.2 | 0.4 |
| *Lhr*<sup>KO</sup>, m | 12 | 1,804 | 15 | 9 | 32 | 2 | 9 | 1 | 24 | 6 | 5 | 12 | 15 | 8 | 0 | 7 | 5 | 7.7 | 0.4 |
| *Lhr*<sup>KO/+</sup>, f | 7 | 740 | 2 | 0 | 0 | 0 | - | 0 | 1 | 0 | - | 0 | 0 | - | 0 | 0 | 0 | 0.4 |  |
| *Lhr*<sup>KO/+</sup>, m | 8 | 871 | 1 | 0 | 0 | 0 | 0 | 0 | 0 | 0 | 0 | 0 | 0 | 0 | 0 | 0 | 0 | 0.1 | 0 |
| Double, f | 20 | 2,364 | 11 | 17 | 30 | 7 | - | 2 | 26 | 3 | - | 20 | 18 | - | 2 | 14 | 5 | 5.7 | 0.6 |
| Double, m | 23 | 2,720 | 17 | 11 | 25 | 4 | 8 | 3 | 22 | 1 | 10 | 23 | 29 | 5 | 2 | 18 | 6 | 5.9 | 0.7 |

CABs were detected in colchicine-treated and hypotonically swollen metaphases of both females (f) and males (m). CD, chromatid deletions (breaks involving a single chromatid). ISOBs isochromatid breaks (both sister chromatids broken at the same location); F, acentric fragment (in ISOBs with F, both the centric and the acentric fragment are present; in ISOBs with no F, only the centric fragment is present; the extra F class includes cells with a complete chromosomal complement accompanied by an extra F). Ah, broken within autosomal heterochromatin; E, chromatid or chromosome exchange; TF, telomeric fusion; H, hyperploid metaphases. Double, *Hmr*<sup>3</sup>; *Lhr*<sup>KO</sup> double mutant. The frequencies of CABs have been calculated without taking into account cells with TFs and hyperploid cells; TF frequencies have been calculated without taking into account hyperploid cells.

**Table 2.** Distribution of heterochromatic breaks among the major autosomes. (observations were limited to metaphases allowing unambiguous recognition of the second and third chromosome. f, females; m, males)

| Mutant, sex | Total breaks | 2<sup>nd</sup> chromosome heterochromatin | 3<sup>rd</sup> chromosome heterochromatin |
| --- | --- | --- | --- |
| *Hmr*<sup>3</sup>, f and m | 47 | 27 | 20 |
| *Lhr*<sup>KO</sup>, f and m | 63 | 38 | 25 |
| *Hmr*<sup>3</sup>; *Lhr*<sup>KO</sup>, f and m | 65 | 32 | 33 |

### Mutations in Hmr and Lhr cause chromosome aberrations but not aneuploid cells in larval brains

To investigate the mitotic roles of *Hmr* and *Lhr* we examined third-instar larval brains from *Hmr*^*3*^ and *Lhr*^*KO*^ homozygotes and from *Hmr*^*3*^*; Lhr*^*KO*^ double mutants. The double mutant was just as viable as the single mutants and did not show any appreciable morphological phenotype. This observation suggests that the simultaneous loss of both Hmr and Lhr is equivalent to the loss of either single protein, consistent with the finding that the Hmr and Lhr proteins are mutually dependent for their stability (Thomae et al., 2013). We incubated brains in saline with colchicine for 1 hour before hypotonic treatment and fixation. Colchicine arrests mitotic cells in metaphase and hypotonic treatment results in chromosome spreading. Preparations obtained in this way allow unambiguous assessment of both chromosome aberrations and telomeric fusions (Gatti and Goldberg 1991; Cenci et al, 1997). Examination of these preparations also allows detection of aneuploid cells. We classified as aneuploid cells only hyperploid figures showing a normal chromosome complement plus an additional unbroken chromosome, all displaying the same degree of mitotic condensation; we did not consider hypoploid cells missing one or more chromosomes because they can occasionally be generated artifactually by the squashing procedure.

Examination of larval brain preparations form *Hmr*^*3*^ and *Lhr*^*KO*^ homozygous larvae and from *Hmr*^*3*^*; Lhr*^*KO*^ double homozygous mutants revealed very similar patterns and frequencies of chromosome abnormalities (Figure 2; Table 1). In single and double mutants ∼5-8% of colchicine-arrested metaphases showed chromosome aberrations, and telomeric fusions were found in ∼0.4-0.9% of metaphases. The frequency of chromosome aberrations in these mutants is more than tenfold higher than that observed in *Hmr*^*3*^*/+* and *Lhr*^*KO*^*/+* heterozygotes or previously observed in Oregon-R controls, all of which showed from 0.3 to 0.7 % cells with chromosome aberrations (Gatti et al., 1974; Gatti, 1979; Benna et al, 2010; Marzio et al 2014; Merigliano et al, 2017). The TF frequencies observed in *Hmr*^*3*^ and *Lhr*^*KO*^ mutants are very low but are nonetheless a clear departure from normality, given that the TF frequency in control cells is virtually zero (Table 1).

Most chromosome aberrations observed in the mutants were isochromatid breaks in which both sister chromatids are broken at the same location. Notably, more than 60% of these isochromatid breaks were incomplete; that is, they consisted either of a centric fragment without the corresponding acentric element or of an acentric fragment associated with a normal chromosome complement (Figure 2). These incomplete isochromatid breaks are rather rare in other mutants that exhibit chromosome aberrations (*mei-9*, *mei-41* (ATR), *mus-102*, *mus-105*, *mus-109*, *tim2*, *dPdxk*, *tws*), where they ranged from 2 to 5% of total isochromatid breaks (Gatti, 1979; Benna et al., 2010; Marzio et al 2014; Merigliano et al., 2017). Such incomplete isochromatid breaks are in contrast very frequent in *Topoisomerase2* (*Top2*) mutants, where they are caused by the rupture of chromosome bridges generated during anaphase due to failure of sister chromatid decatenation (Mengoli et al. 2014). Although incomplete isochromatid breaks and at least some complete isochromatid breaks are likely to result from breaks generated during anaphase (Figure 3), chromatid deletions (breaks involving only one of the two sister chromatids; see Figure 2 and Table 1) cannot be the outcome of anaphase defects but rather must result from lesions produced during S or G2 phase. Thus, the low frequencies of chromatid deletions observed in *Lhr* and *Hmr* (∼0.1-0.8%) are likely to reflect the presence of double strand breaks leading to chromosome breakage (Obe et al. 2002; Durante et al. 2013).

**Figure 3.**
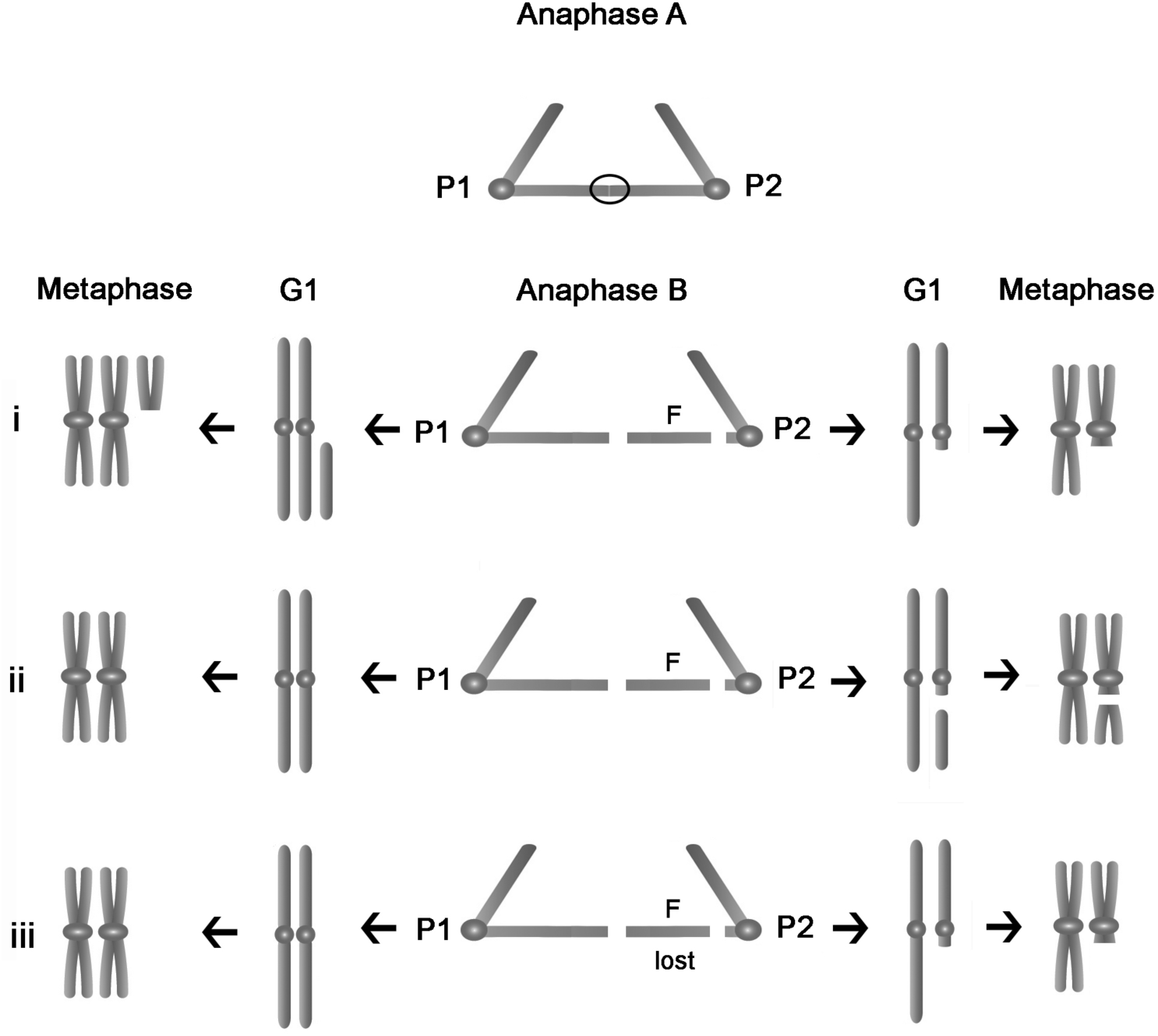
A model for the formation of incomplete isochromatid breaks (ISOBs). The primary event leading to an ISOB is the formation of a chromatin bridge generated by transient sister chromatid association during anaphase A, represented by an ellipse (see Discussion for possible origin of this association). Anaphase A is characterized by sister chromatid separation and poleward movement of the daughter chromosome sets. In anaphase B the spindle poles move apart, increasing the distance between the two daughter chromosome sets. The anaphase drawing refers to the mitosis in which the anaphase bridge forms and depicts a single chromosome composed of a pair of sister chromatids; centromeres are represented by circles. G1 and metaphase refer to the subsequent cell cycle and show both homologous chromosomes, one of which segregated normally during the previous anaphase. The metaphase configurations are those observed in colchicine treated cells of mutants that are shown in Figure 2. Stretching of the bridge during anaphase B would result in the resolution of the sister chromatid association and a rupture at the heterochromatin-euchromatin junction. This situation could have three possible outcomes: (i) The acentric fragment (F) segregates with the chromatid to which it was originally attached giving rise to a telophase nucleus containing both homologous chromosomes plus an acentric element (at cell pole 1, P1), and to another nucleus containing only the centric element and no corresponding acentric fragment (at cell pole 2, P2). (ii) The acentric fragment segregates with its corresponding centric element producing a normal nucleus at P1 and a nucleus bearing a complete ISOB at P2. (iii) The acentric fragment is lost leading to a normal telophase nucleus at P1 and to a nucleus containing a broken chromosome without the corresponding fragment at P2. This model describes the possible outcomes of a rupture at the heterochromatin-euchromatin junction but could also be extended to ruptures in the euchromatin.

In both *Hmr* and *Lhr* mutants, more than 70% of all breaks are located in heterochromatin or at the junction between euchromatin and heterochromatin (henceforth designated as heterochromatic breaks). In previous studies, mutants in *mei-9*, *mei-41*, *mus-102* and X-ray treated wild type cells displayed 40-50% heterochromatic breaks, while mutants in *mus-105* and *mus-109* showed 18% and 81% heterochromatic breaks, respectively (Table S1). The proportion of breaks in the Y chromosome in males is also higher for *Hmr* and *Lhr* than any previous condition analyzed. It is likely that the high frequency of heterochromatic breaks observed in *Hmr* and *Lhr* mutants reflects a specific fragility of heterochromatin and euchromatin-heterochromatin junctions during anaphase, similar to that observed in *Top2* mutants (Mengoli et al, 2014). However, while in *Top2* mutants most incomplete isochromatid breaks involve the Y chromosome and a distal heterochromatic region in the 3L arm (region h47), in *Hmr* and *Lhr* mutants isochromatid breaks involve the Y and both the second and the third chromosome heterochromatin (Tables 1 and 2). Assessing isochromatid breaks within the X chromosome heterochromatin was difficult because breaks that separate the DAPI-bright from DAPI-dull region of X heterochromatin produce fragments that closely resemble a fourth chromosome and an autosomal arm, respectively.

In both the single and the double mutants, hyperploid cells were quite rare, ranging from only 0.2 to 0.3%. This is not due to a low survival rate or low division potential of hyperploid cells, because this type of cell is very frequent in *Drosophila* mutants defective in chromosome segregation. For example, mutants in the *zw10* gene that encodes a component of the spindle checkpoint machinery display 50-60% hyperploid cells (Smith et al., 1985; Williams et al., 1992). Similarly, mutants in the *mitch* gene that specifies a subunit of the Ndc80 kinetochore complex exhibit 43% hyperploid cells (Williams at al., 2007).

### Analysis of cell division in non-colchicine-treated cells from Hmr and Lhr mutant brains reveals abnormal anaphases

To obtain insight into the mechanism leading to the incomplete chromosome aberrations observed in colchicine treated cells, we examined mitotic division in mutant brains in the absence of colchicine or hypotonic treatment, in order to directly visualize anaphase. We first determined the mitotic index and the frequencies of anaphases in *Hmr*^*3*^*; Lhr*^*KO*^ double mutants, in which the function of the Hmr-Lhr complex should be eliminated completely. *Hmr*^*3*^*; Lhr*^*KO*^ brains displayed a mitotic index and a frequency of anaphases comparable to those observed in wild type controls, suggesting that mutant cells progress through mitosis at the same rate as wild type cells (Table 3).

**Table 3.** *Hmr; Lhr* double mutants exhibit normal mitotic parameters. (Brains were fixed without colchicine or hypotonic treatment, squashed and stained with DAPI to visualize chromosomes. f, females; m, males)

| Genotype, sex | # of fields | Prophases and prometaphases | Metaphases | Anaphases | Divisions/ fields | Anaphases (%) |
| --- | --- | --- | --- | --- | --- | --- |
| wt control, f | 180 | 292 | 40 | 72 | 2.2 | 14.8 |
| wt control, m | 172 | 304 | 54 | 62 | 2.4 | 17.8 |
| *Hmr*<sup>3</sup>; *Lhr*<sup>KO</sup>, f | 338 | 720 | 103 | 177 | 3.0 | 17.7 |
| *Hmr*<sup>3</sup>; *Lhr*<sup>KO</sup>, m | 261 | 540 | 68 | 116 | 2.8 | 16.0 |

We next examined anaphase figures in both *Hmr* and *Lhr* single mutants and *Hmr; Lhr*, double mutants. We found that a substantial fraction of mutant anaphases (ranging from ∼12 to 16.5%; Figure 4, Table 4) display chromosome bridges, bridges plus fragments, or acentric fragments, but no anaphases showed intact lagging chromosomes. These observations support the hypothesis that the incomplete aberrations shown in Figure 2 were generated by chromosome breakage occurring during a previous anaphase. The defective anaphases observed in *Hmr* and *Lhr* mutants are unlikely to be the outcome of telomeric fusions, as the TF frequency is approximately 20-fold lower than that of aberrant anaphases (Table1).

**Table 4.** Anaphase defects in *Hmr* and *Lhr* mutants.

| Genotype, sex | # of brains | # of anaphases | Bridge or broken bridge | Bridge and fragment | Acentric fragment | % defective |
| --- | --- | --- | --- | --- | --- | --- |
| wt control, f | 7 | 104 | 1 | 0 | 1 | 1.9 |
| wt control, m | 8 | 115 | 2 | 0 | 0 | 1.7 |
| *Hmr*<sup>3</sup>, f | 9 | 78 | 4 | 4 | 2 | 12.8 |
| *Hmr*<sup>3</sup>, m | 9 | 116 | 7 | 4 | 6 | 14.7 |
| *Lhr*<sup>KO</sup>, f | 8 | 101 | 8 | 2 | 2 | 11.9 |
| *Lhr*<sup>KO</sup>, m | 19 | 326 | 29 | 10 | 7 | 14.1 |
| *Lhr*<sup>KO</sup>; *Hmr*<sup>3</sup>, f | 13 | 286 | 16 | 15 | 8 | 13.6 |
| *Lhr*<sup>KO</sup>; *Hmr*<sup>3</sup>, m | 11 | 237 | 19 | 14 | 6 | 16.5 |

f, females; m, males

**Figure 4.**
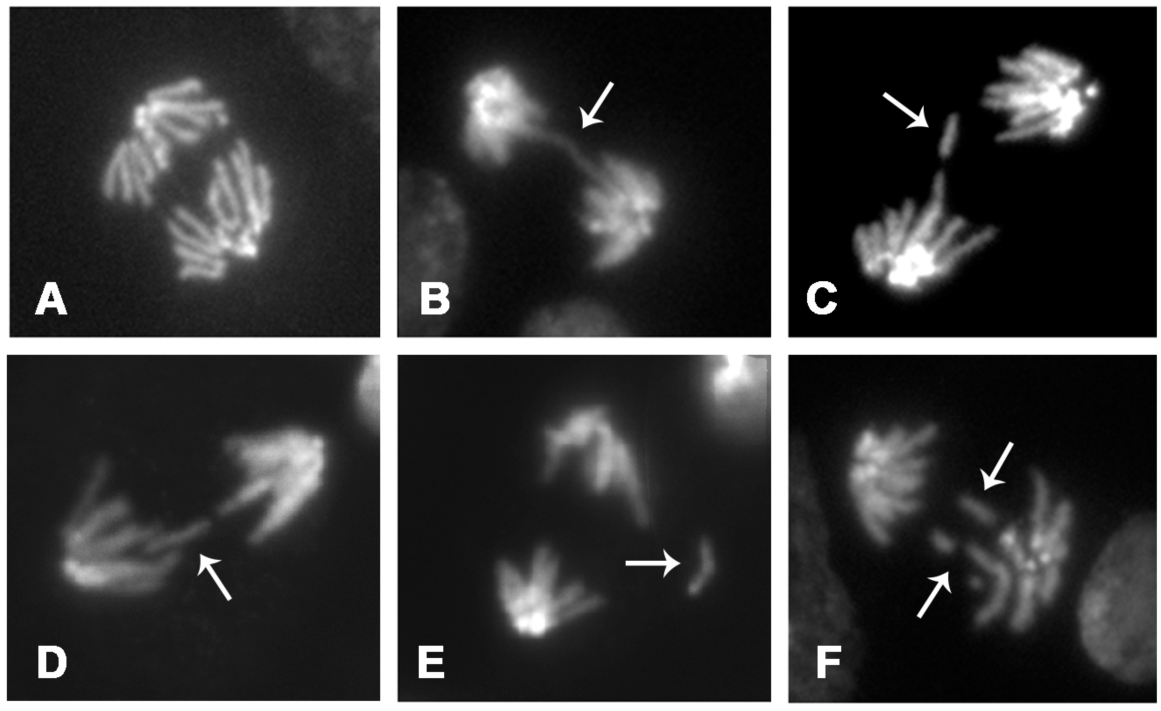
Examples of aberrant anaphases observed in *Lhr* and *Hmr* mutants. A. Wild type control. B. Anaphase with bridge. C and D. Anaphases with a broken bridge and a fragment. E. Anaphase with a fragment. F. Anaphase with fragments.

Finally, we examined preparations fixed with formaldehyde and stained for tubulin and DNA, and counted the cells with prometaphase/metaphase spindles showing aligned or unaligned chromosomes. This analysis revealed no differences between wild type controls and mutant cells, suggesting that neither *Hmr* nor *Lhr* are required for formation of the metaphase plate (Figure 5; Table 5).

**Table 5.** *Hmr* and *Lhr* mutants exhibit normal chromosome alignment at metaphase.

| Genotype, sex | # of prometaphases* and metaphases | Non-aligned | Aligned | Aligned (%) |
| --- | --- | --- | --- | --- |
| wt, control, f + m | 167 | 70 | 97 | 58 |
| *Hmr*<sup>3</sup>, f + m | 226 | 104 | 122 | 54 |
| *Lhr*<sup>KO</sup>, f + m | 256 | 103 | 153 | 60 |
| *Hmr*<sup>3</sup>; *Lhr*<sup>KO</sup>, f + m | 182 | 82 | 100 | 55 |

\* We considered only prometaphases with fully formed spindles (similar to metaphase spindles) with chromosomes showing the same degree of condensation as metaphase chromosomes. f, females; m, males.

**Figure 5.**
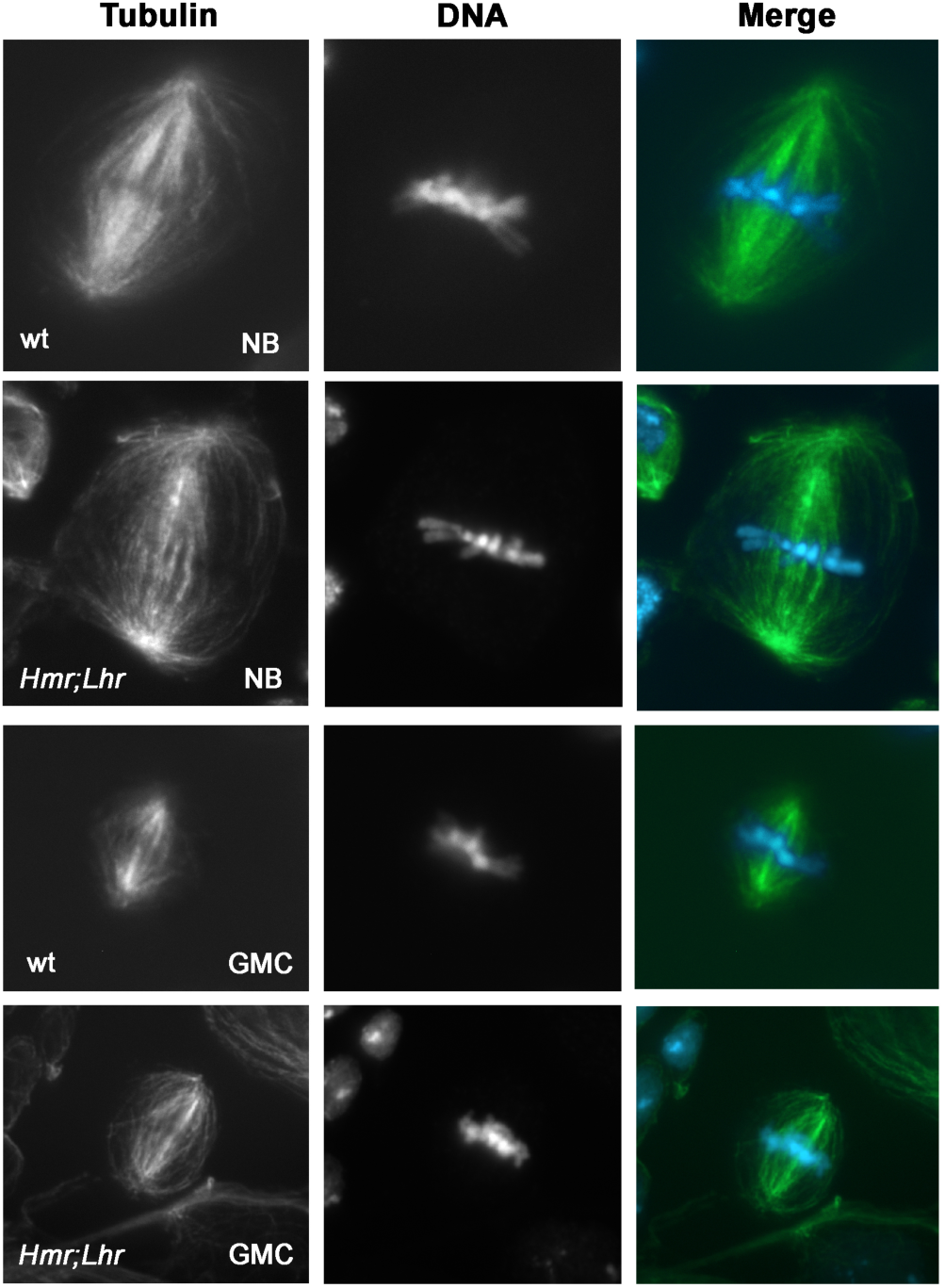
Examples of well-aligned metaphases observed in brains from wild type (wt, Oregon R) and *Hmr; Lhr* double mutants. NB, neuroblast; GMC, ganglion mother cells.

### Dynamic behavior of Hmr and Lhr during mitotic division

To gain further insight into the roles of Hmr and Lhr, we stained brain preparations from larvae that express either Hmr-HA or Lhr-HA with anti-HA and anti-Cid (a centromere marker homologous to Cenp-A) antibodies. We found that Hmr and Lhr exhibit very similar dynamic behaviors in both male (Figures 6 and 7) and female cells (Figures S1 and S2). In interphase cells, Lhr and Hmr showed similar localizations near but not overlapping with the DAPI-bright heterochromatin and did not co-localize with Cid (Figures 6A, 7A, S1A and S2A). This localization pattern suggests that both proteins are enriched in heterochromatin regions that are not fluorescent after DAPI or Hoechst 33258 staining (see Gatti and Pimpinelli, 1992, for a map of mitotic heterochromatin). In the only very early prophase we were able to find, where only heterochromatin has started to condense and euchromatin is still diffuse, Lhr was clearly associated with heterochromatin and concentrated in chromosomal regions that are not DAPI-bright (Figure 7B). Lhr was also associated with dully-fluorescent heterochromatic regions in another early prophase in which euchromatin was visible but poorly condensed (Figure S2B). In all other prophases stained for Lhr (9 male and 10 female prophases), Lhr was not associated with the chromosomes and exhibited a diffuse nucleoplasmic localization (Figures 7C, and S2C). Hmr was dispersed within the nucleoplasm and did not exhibit a clear accumulation in dully-fluorescent heterochromatic regions of the chromosomes in all the 5 male and 7 female prophases scored. We only observed an occasional Hmr accumulation at a small pericentromeric chromosomal region (figure S1B) (likely 2R heterochromatin, based on its localization on metaphase chromosomes). However, none of the prophases stained for Hmr was a very early one, like that shown for Lhr in Figure 7C. Thus, we cannot exclude that Hmr remains associated with heterochromatin during the earliest stages of prophase like Lhr does. Regardless of this possible small difference in behavior, however, our results strongly suggest that both Lhr and Hmr mostly dissociate from the chromosomes during early prophase.

**Figure 6.**
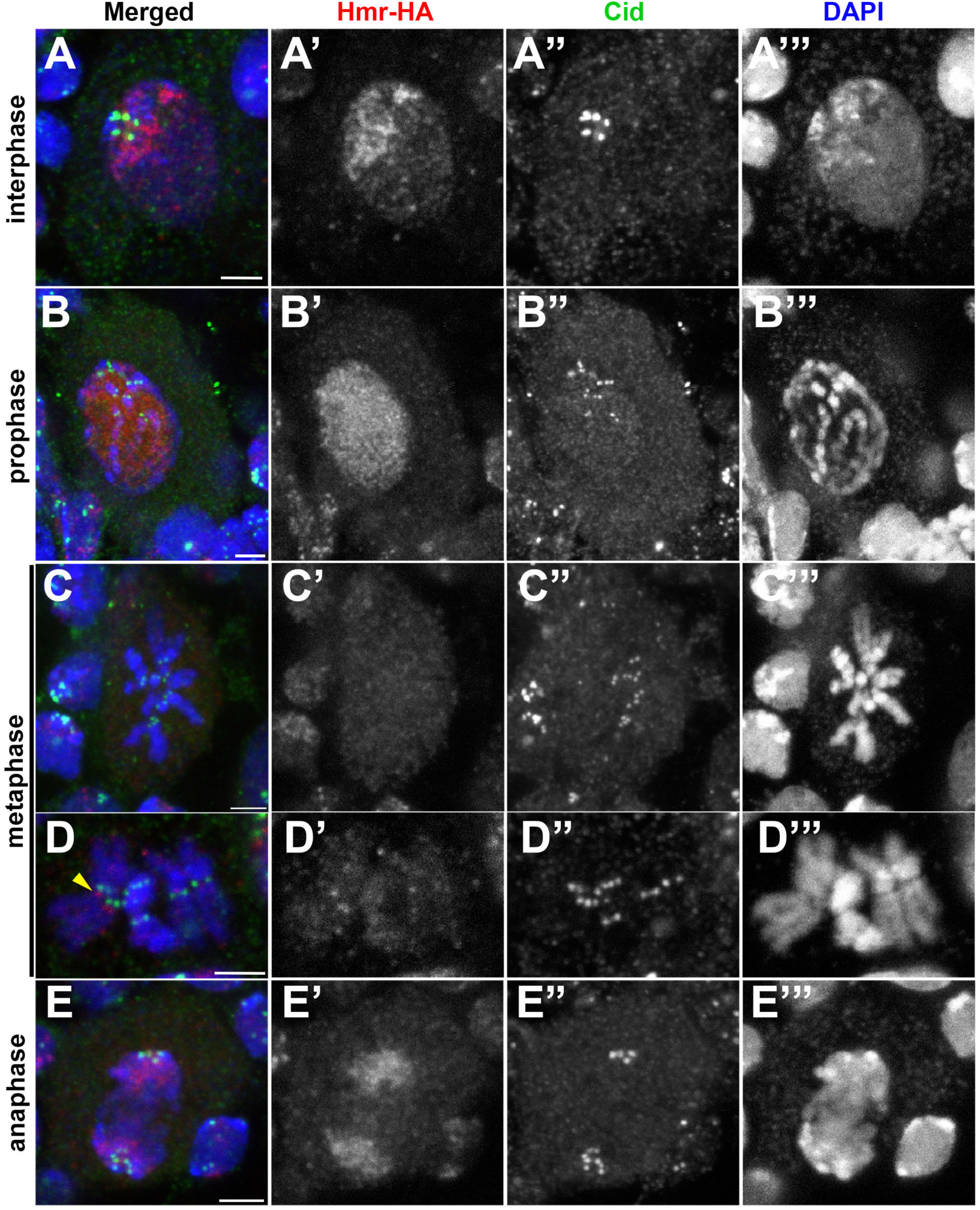
Localization of Hmr in male brain cells. Note the dynamic behavior of the bulk of the protein; in interphase, it is nuclear and does not colocalize with the Cid signals (A), it dissociates from the chromosomes during prophase (B) and metaphase (C and D) and reassociates with the chromosomes during anaphase (E). A fraction of metaphases showed Hmr accumulations on the 2R heterochromatin (arrowhead) (D). See text for a detailed description of the dynamic localization of Hmr.

**Figure 7.**
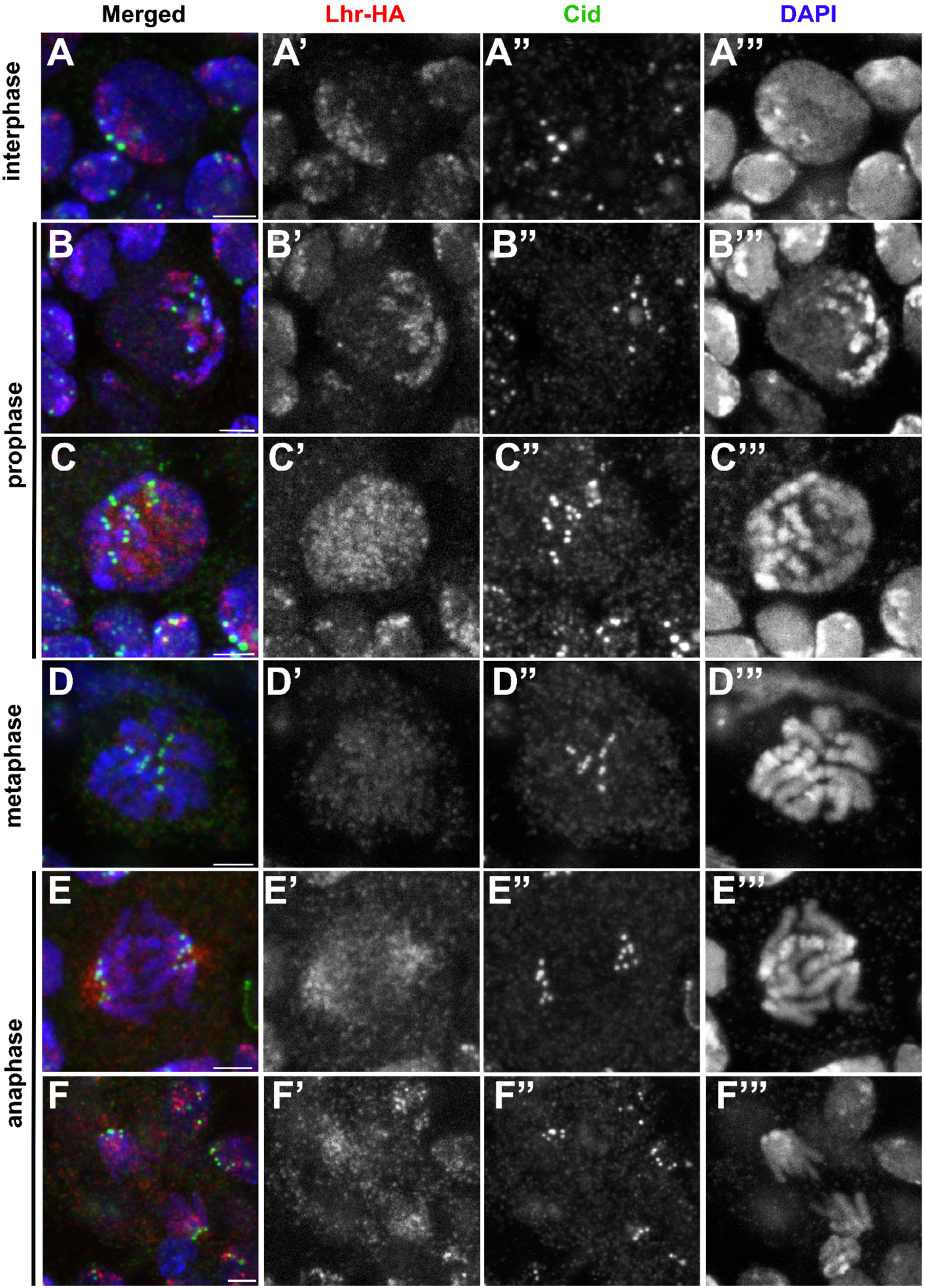
Localization of Lhr in male brain cells. Lhr does not colocalize with the Cid signals in interphase nuclei (A). Note that Lhr exhibits a dynamic behavior similar to that of Hmr (shown in figure 6). In very early prophase, in which only heterochromatin is condensed, Lhr is still associated with the heterochromatin (B), but dissociates from the chromosomes in both late prophase (C) and metaphase (D) cells. Also note that in the early anaphase shown (E), Lhr localizes in regions distal to the Cid signals that probably correspond to the spindle poles; in the late anaphase (F) Lhr is instead localized proximally to the spindle signals and is incorporated into the heterochromatin. See text for a detailed description of the dynamic localization of Lhr.

Lhr was consistently dissociated from the chromosomes in prometaphase (n=14) and metaphase (n=11) cells (Figures7D and S2D). Hmr too was mostly dissociated from prometaphase and metaphase chromosomes (Figures 6C, S1D), but in a fraction of the cells (4/12 in males, 12/28 in females) a small amount of Hmr was concentrated in a single non-fluorescent region of the 2R heterochromatin (region h42 according to the heterochromatin map of Gatti and Pimpinelli, 1992), well separated from the centromeric region marked by Cid (Figures 6D and S1C). Because this accumulation of Hmr in region h42 is seen in prophase (Figure S1B), it likely results from a failure of Hmr to dissociate from heterochromatin when cells enter mitosis. During anaphase, both Hmr and Lhr become incorporated again into heterochromatin (figures 6E, n=6; 7E, F, n=7; S1E, n=11; S2E, n=3). In some early anaphases, Lhr was concentrated in regions distal to both the chromosomes and the Cid signals, which likely to correspond to the spindle poles (Figure 7E). This observation raises the intriguing possibility that Lhr travels towards the spindle poles along the microtubules, so as to favor its recruitment by the pericentric heterochromatin. Finally, we note that in both Hmr-HA and Lhr-HA expressing cells, the non-chromocentral regions of interphase nuclei were faintly stained, suggesting that while the bulk of Hmr and Lhr is in the heterochromatin, small amounts of these proteins are associated with euchromatin.

In summary, these results show that Lhr and Hmr colocalize with the non-DAPI-bright heterochromatin but not with the Cid-stained centromeres during interphase. Both Lhr and Hmr mostly dissociate from heterochromatin during prophase, and return to the heterochromatin during anaphase. To explain the slightly different dynamic behaviors of Lhr and Hmr we hypothesize that the two proteins usually form a complex when incorporated into heterochromatin but are partially independent when they dissociate and reassociate with heterochromatin. The results reported here agree with previous observations in embryos, showing that Lhr does not co-localize with Cid signals or with the 359 bp and the AATAT satellite DNAs, which are both strongly fluorescent after DAPI staining (Maheshwari and Barbash, 2012). However, Maheshwari and Barbash (2012) showed that in embryonic metaphases Lhr concentrates next to the dodecasatellite DNA that marks the third chromosome, while here we observed Hmr but not Lhr localization to chromosome 2 heterochromatin. These results are subject to two interpretations. It is possible that Lhr concentrates in the third chromosome heterochromatin also in brain cells, and that we failed to detect it due to differences in the fixation and/or immunostaining procedures. Alternatively, Lhr might specifically accumulate in the third chromosome heterochromatin only in embryonic cells. Regardless, the different chromosomal localizations during metaphase reinforce the conclusion that Hmr and Lhr can have partially independent localizations.

### Aberrant chromosome condensation but normal centromere function in hybrid males

We next turned to an analysis of the mitotic phenotype of brains from hybrid third-instar larvae generated by crosses between *D. melanogaster* females and *D. simulans* males, in order to determine whether or not hybrids show similar phenotypes to the *D. melanogaster Hmr* and *Lhr* mutants. We first examined colchicine-treated metaphases from larval brains of viable hybrid females, and found that the chromosomes of these metaphases are morphologically indistinguishable from those of wild type females. In addition, we found that larval brain metaphases of these females have a very low rate of chromosome aberrations (<1%; Table S2), fully comparable to that of wild type controls.

Consistent with previous results (Orr et al., 1997; Bolkan et al., 2007), hybrid male larvae displayed small brains, were devoid of imaginal discs, and had low frequencies of mitotic divisions; hybrid male cells were found to be predominantly stalled in interphase with only a few M-phase cells escaping the interphase block (Bolkan et al., 2007). To define the mitotic phenotype in these hybrid brains we first examined brain squash preparations, without colchicine and hypotonic pretreatments. This analysis revealed that hybrid male brains exhibit an about 5-fold reduction of the mitotic index compared to brains from either *D. melanogaster* or *D. simulans* third instar larvae, a reduction also seen previously (Bolkan et al., 2007).

To evaluate chromosome condensation and integrity, we next treated hybrid male brains with hypotonic solution in the absence of colchicine pretreatment, and then immunostained preparations with an anti-phospho histone H3 antibody that marks mitotic chromatin (Wei et al., 1999). We found several types of defects in chromosome structure, and none of the 100 cells we examined appeared completely normal. 31% of mitotic cells were prophase-like cells with elongated and poorly condensed chromosomes enriched in phospho histone H3 (Figure 8B). Similar cells were also observed in third-instar larval brains of *D. melanogaster* and *D. simulans* males, but in each species prophases accounted for no more than 10% of the mitotic figures (*n* > 100 in each species). An increase in prophase cells in hybrids was previously observed by Bolkan et al., (2007). This finding suggests that in hybrid male prophases, chromosome condensation is delayed compared to the parent species. The hybrid male brains also showed substantial defects in chromosome condensation at later mitotic stages. 50% of dividing cells were prometaphase/metaphase figures with fuzzy chromosomes in which the sister chromatids were closely apposed; in addition, these cells often displayed obvious chromosome breaks (Figure 8C and Figure S3). 15% of the mitotic cells in hybrid male brains were prometaphases/metaphases with well-separated sister chromatids, but in these cells the chromosomes were fuzzy and overcondensed compared with those of the parent species. Although the hypotonic treatment reduces the frequency of anaphases (Gatti and Baker, 1989), we were able to observe 4 anaphases (4% of the mitotic figures), which displayed a higher degree of chromosome condensation than seen in non-hybrid larvae from either species (Figure 8D).

**Figure 8.**
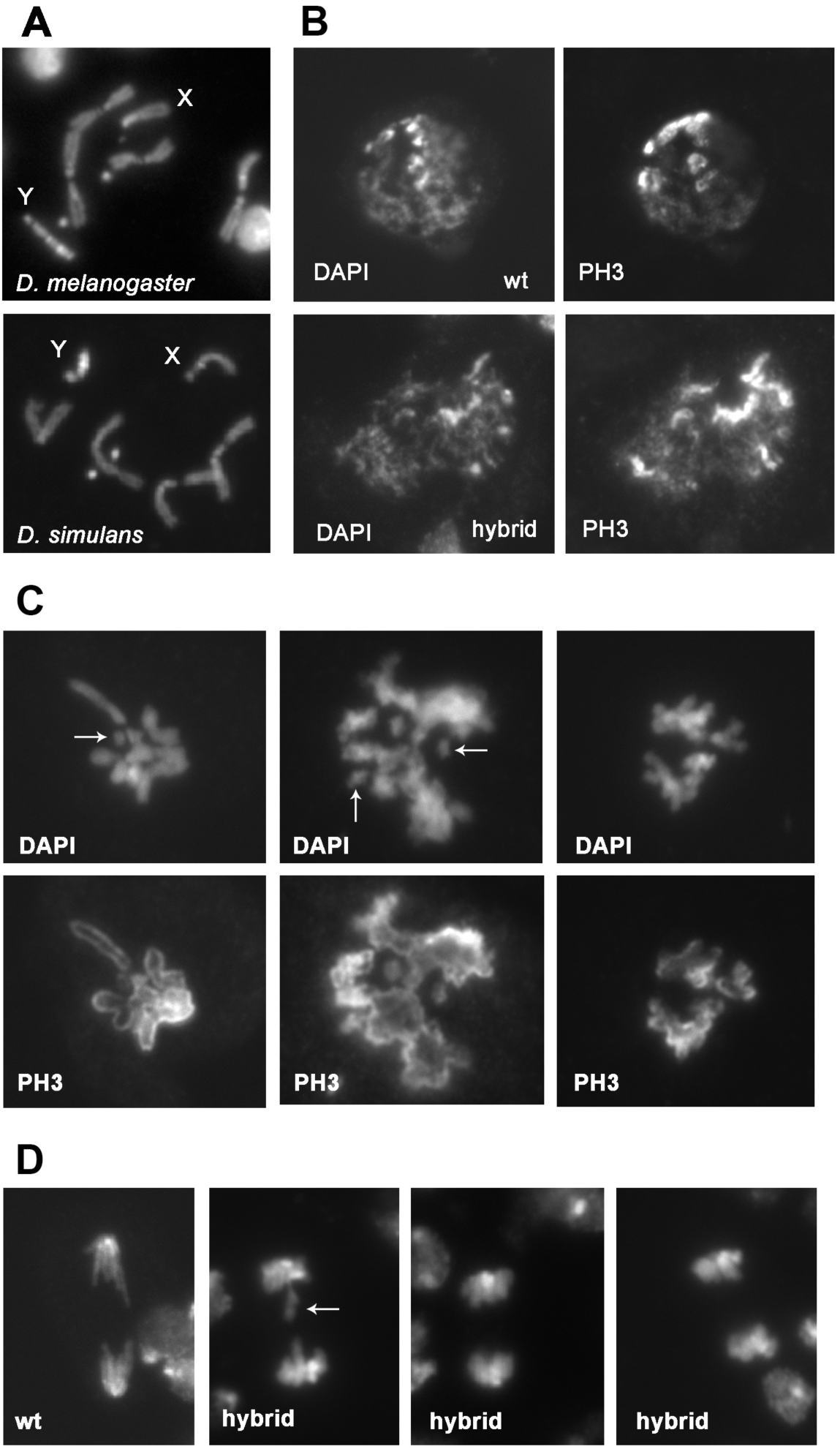
Hybrid males exhibit defects in chromosome condensation and integrity. Larval brains were fixed with formaldehyde but not treated with colchicine or hypotonically swollen (see Methods), and then stained for DNA (DAPI) and the mitotic phospho histone H3 (PH3). A. Male prometaphases/metaphases from wild type *D. melanogaster* and *D.* simulans; note the differences in the X chromosome heterochromatin staining and in the size of the Y chromosome. B. Prophase-like figures from wild type *D. melanogaster* and hybrid males. C. Prometaphases/metaphases from hybrid males with poorly condensed and broken (arrows) chromosomes. Note that in two of the cells shown the sister chromatids are fuzzy and closely apposed. D. Anaphases observed in wild type *D. melanogaster* and in hybrid males; note that the anaphase chromosomes in the hybrids are more condensed than in control; one of them also exhibits a broken bridge (arrow).

To analyze mitotic divisions in a broader context, hybrid male brains not exposed to hypotonic treatment were stained for DNA and with anti-tubulin antibodies. Dividing cells in these brains indeed form a mitotic spindle. Of the mitotic figures scored (52 excluding prophase-like figures, from 12 brains), 37% were prometaphases, 52% metaphases with well-aligned chromosomes, and 11% anaphases (Figure 9). In wild type *D. melanogaster* male brains, these mitotic figures (*n* = 100) were 36%, 44% and 20% respectively; in *D. simulans* (n= 89) they were 30%, 49%, and 21%, respectively. Thus, the rare dividing cells of hybrid males exhibit clear defects in chromosome condensation and integrity. However, these cells appear to have a normal centromere/kinetochore function, as suggested by the ability of the chromosomes to congregate in a metaphase plate and undergo anaphase.

**Figure 9.**
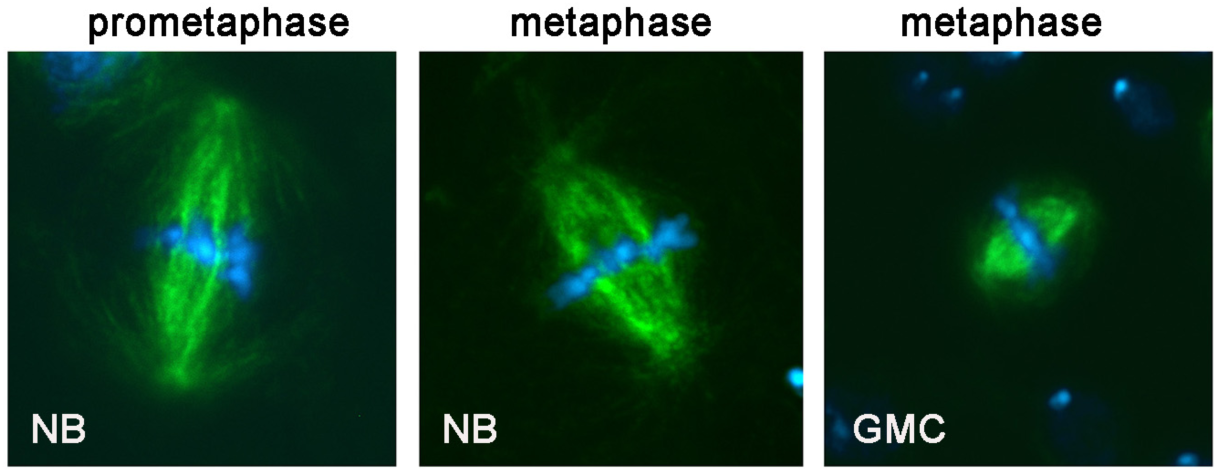
Hybrid males form a mitotic spindle and do not exhibit defects in chromosome alignment at metaphase. A prometaphase and two well-aligned metaphases observed in hybrid males. For control metaphases see Figure 5; NB, neuroblast; GMC, ganglion mother cell.

## Discussion

### Hmr and Lhr do not have a major role in centromere function

Our data clearly show that *Hmr* and *Lhr* mutants and *Hmr*; *Lhr* double mutants exhibit very similar patterns of chromosome abnormalities. In none of the mutants did we observe hyperploid cells, which are very common in mutants that disrupt kinetochore function or the spindle checkpoint machinery (Smith et al., 1985; Williams et al., 1992; Williams et al., 2007). Furthermore, we found that the mutants exhibit mitotic indexes and frequencies of well-aligned metaphase plates comparable to those seen in wild type controls. However, we discovered that the mutants have ∼6% chromosome aberrations, most of which were incomplete isochromatid breaks that are likely to be generated during anaphase (Mengoli et al., 2014). Consistent with this idea, 15% of anaphases showed chromatin bridges, broken bridges, and/or acentric fragments caused by breaks in heterochromatin or at the euchromatin/heterochromatin junction. Importantly, however, we never observed anaphases with intact lagging chromosomes, which indicates that the centromeres are properly aligning during metaphase and separating during anaphase. All mutants also showed low frequencies of telomeric fusions. Thus, *Lhr* and *Hmr* mutants have a normal centromere and kinetochore function and a weak deficiency in telomere capping but are defective in the separation of sister chromatids during anaphase.

Previous work described Hmr and Lhr as centromere proteins, as reviewed here in the Introduction (Thomae et al., 2013). We analyzed the localization of Hmr-HA and Lhr-HA in larval brains and were not able to confirm their centromeric localization. In interphase nuclei Hmr-HA and Lhr-HA were enriched in the chromocenter but did not colocalize with the Cid signals, in contrast to Thomae et al. (2013). In metaphase, Hmr- and Lhr were also not accumulated on the centromeres, consistent with the observations of Thomae et al. (2013), though we did find that Hmr is enriched in a region of the 2R hetrochromatin. In addition, we were not able to confirm the major cytogenetic evidence reported by Thomae et al. (2013) for a defect in centromere function in Hmr- and Lhr-depleted cells, namely the presence of lagging chromosomes in anaphases. Although we also found lagging chromatin in mutant anaphases, Figures 2 and 4 clearly demonstrate that this material consists of chromosome fragments rather than intact chromosomes. In addition, we did not observe hyperploid cells in the mutants, but instead mostly cells with incomplete isochromatid breaks. Thus, our data argue strongly against a major functional role of Hmr and Lhr at *Drosophila* centromeres or kinetochores.

### Hmr and Lhr have a role in sister chromatid separation during anaphase

One of the main questions raised by our results is the mechanism leading to the formation of aberrant anaphases in mutant cells. The main defects observed in the anaphases are continuous or broken bridges between the two daughter chromosome sets. In theory, these chromatin bridges and their broken derivatives might originate from telomeric fusions involving either sister chromatids or single non-sister chromatids. However, bridge formation from such telomeric fusions is unlikely because (i) unambiguous sister telomere fusions were never observed in either wild type and mutant brains, and (ii) fusions between non-sister telomeres, which were present in less than 1% of the cells, cannot account for ∼15% aberrant anaphases (Tables 1 and 3). Another possible source of anaphase bridges are isochromatid breaks with fusion of the proximal broken ends (sister union). The presence of a proximal sister union can only be detected in isochromatid breaks broken in euchromatin; if breaks occurred at the euchromatin-heterochromatin junction or purely in heterochromatin, the close apposition of the proximal broken ends prevents reliable evaluation of their possible fusion. Two reasons, however, lead us to believe that the bridges we observe are not generated by sister union isochromatid breaks. First, we did not observe clear sister unions in the few euchromatic isochromatid breaks we detected. Second, sister union heterochromatic isochromatid breaks are expected to give rise to very short bridges and not to the long bridges we observed in the aberrant anaphases.

The most likely mechanism for the anaphase bridges we observed involves a defect in sister chromatid separation and not a centromere or kinetochore dysfunction (Figure 3). Sister chromatid separation is a complex process. First, to segregate properly, sister chromatids must condense through the action of the condensin complexes. Indeed, mutations in the *Drosophila* condensin subunit coding genes *gluon* (SMC4) and *barren* (XCAP-H) result in abnormally condensed chromosomes and formation of extensive chromatin bridges at anaphase (Bhat et al., 1996; Steffensen et al., 2001; Somma et al., 2003). Second, the DNA molecules of the two sister chromatids must be decatenated through the action of topoisomerase II. Accordingly, anaphase chromatin bridges have been observed in *Drosophila* S2 cells depleted of Topoisomerase II (Chang et al, 2003; Somma et al., 2008; Coelho et al., 2008) and in *Top2* mutant brains (Mengoli et al. 2014). Some investigators have posited a relationship between these two steps, suggesting that condensin is required for the organization of a correct axial chromatid structure in which topoisomerase II can efficiently promote sister chromatid decatenation (Coelho et al., 2003). Third, separase mediates the proteolytic degradation of the cohesin complex, so as to allow sister chromatid separation at the metaphase-to anaphase transition. In interphase cells of both *Drosophila* and vertebrates, cohesin localizes to both heterochromatin and euchromatic chromosome arms. However, the bulk of cohesin is released from the chromosomes during prophase through a separase-independent mechanism, and returns to the chromosomes during telophase; only very small amounts of cohesin persist between the sister chromatids during prometaphase and metaphase (reviewed by Dorset and Ström, 2012; Hirano, 2015; Uhlmann, 2016). Thus, the cohesins appear to behave like Hmr and Lhr. Notably, the *Drosophila* cohesin-loading factor Nipped-B (NIPBL/Scc2) accumulates in heterochromatin including the ectopic heterochromatin regions generated by translocations. Germane to Hmr and Lhr loss of function phenotype, Nipped-B/cohesin accumulation in these regions delays sister chromatid separation during anaphase, leading to stretched chromatids that bridge the two segregating chromosome sets (Oliveira et al., 2014).

At this stage, we can only speculate about the primary defect in sister chromatid separation leading to chromatin bridges in the mutants. The findings that in *Hmr* and *Lhr* mutants metaphase chromosomes are morphologically normal, and anaphase and metaphase defects are rather rare, suggest that potential variation in condensin or Top2 levels would be minimal and difficult to detect. It is also unlikely that these defects are caused by a direct or indirect defect in Top2 function, because brains from larvae bearing weak mutations in the *Top2* gene exhibit a highly specific pattern of incomplete aberrations, involving specific regions of the Y chromosome and a single region of 3L heterochromatin (region 47) (Mengoli et al., 2014). In contrast, in *Hmr* and *Lhr* mutants we observed breaks in both 3L and 3R heterochromatin and in the second chromosome heterochromatin. Thus, we favor the possibility that the anaphase bridges observed in *Hmr* and *Lhr* mutant cells result from an aberrant sister chromatid cohesion established during interphase that becomes phenotypically manifest when cells enter anaphase.

Regardless of the underlying mechanism, our observations raise the question of why most complete and incomplete isochromatid breaks are broken at the euchromatin-heterochromatin junctions. The transition between heterochromatin and euchromatin appears to be gradual rather than abrupt in terms of DNA content. The junctions do not contain specific DNA sequences and are instead mosaics of middle-repetitive DNA interspersed with single-copy DNA (Miklos and Cotsell, 1990; Hoskins et al., 2002). However, there is an obvious change in DNA fiber compaction at the euchromatin-heterochromatin transition that might render these regions prone to breakage when chromatin bridges are pulled by the spindle. To the best of our knowledge, the literature about preferential sites for anaphase bridge rupture is rather limited. In vivo analysis of human cancer cells containing marked repeated DNA inserted into the chromosomes showed that bridges are preferentially severed at sites near the inserted DNA (Shimizu et al., 2005). In addition, in fixed cells from embryogenic callus cultures, most anaphase bridges occurred within heterochromatic knobs or at the junction between euchromatin and the knobs (Fluminhan and Kameya, 1996). These reports support the hypothesis that euchromatin-heterochromatin transition regions in anaphase bridges are more prone to breakage than euchromatic regions when pulled by spindle-generated forces.

### Do hybrids have a mitotic defect similar to the Hmr and Lhr phenotype in D. melanogaster?

Our results strongly suggest that loss of the Lhr-Hmr complex in *D. melanogaster* results in abnormal adhesion between sister chromatids that leads to anaphase bridges and chromosome breakage. The lethality of interspecific hybrids, however, results from the presence of wild type *Hmr* and *Lhr*, not their loss. We therefore re-examined the chromosomal phenotype of dying hybrid males to investigate whether or not it shows any similarity to *Hmr* and *Lhr D. melanogaster* mutants. The detectable but non-essential roles of *Hmr* and *Lhr* in fundamental functions such as the control of transposition, telomere homeostasis and sister chromatid separation suggest that these genes might play redundant functions in some essential processes, but may or may not offer insight into why mutations in *Hmr* and *Lhr* suppress hybrid male lethality. This uncertainty reflects what may be a general property of hybrid incompatibility genes: Hybrid phenotypes often represent a gain-of-function because they are caused by the presence of wild-type alleles, while the intraspecific phenotypes are typically discovered and analyzed using loss-of-function mutations (Maheshwari and Barbash, 2011).

Previous research showed that most brain cells of hybrid males are arrested in either the G1 or G2 phase and rarely enter mitosis (Bolkan et al. 2007). It has been also reported that a fraction of these cells exhibits highly aberrant chromatin morphology, suggesting that they represent mitotic cells with severely undercondensed chromosomes (Orr et al. 1997). Furthermore, it has been suggested that hybrid male cells have some form of chromatin defect specific to the X chromosome, a hypothesis supported by interactions between dosage compensation mutations and hybrid viability (Barbash, 2010). It is challenging to determine if hybrids show a phenotype at least in part comparable to that seen in *Hmr* and *Lhr* mutants, due to the relatively low penetrance of these mutant phenotypes in *D. melanogaster* and the low frequency of mitotic cells in hybrids. We were able to image sufficient mitotic cells in hybrids to conclude that chromosomes align properly during metaphase. Based on this result and the small number of anaphases observed, there is no evidence that hybrid lethality results from centromere or kinetochore dysfunction (Figure 9). We did though identify defects in chromosome condensation and integrity in hybrid males that are reminiscent of those observed in cells depleted of condensins or showing abnormal accumulations of cohesins (Figure 8; Bhat et al., 1996; Steffensen et al., 2001; Somma et al., 2003; Cobbe et al., 2006; Savvidou et al. 2006; Oliveira et al., 2014). Interestingly, these defects have been shown to interfere with sister chromatid separation leading to anaphase bridges. It will therefore be of interest in future studies to determine whether *Hmr* and *Lhr* mutants and interspecific hybrids have altered distributions and functional defects in condensins, cohesins, or cohesin loading/releasing factors.

## Acknowledgments

J.A.B was the he recipient of a student grant from the U.S.-Italy Fulbright Commission. This work was supported in part by a PRIN/MIUR grant to S. B., a grant from Associazione Italiana per la Ricerca sul Cancro (IG 16020) to M. G., and by NIH R01GM074737 to D.A.B. Y.Y. is supported by the Howard Hughes Medical Institute. We thank Dean Castillo, Anne-Marie Dion-Côté, Michael Goldberg, and Sarah Sander for helpful comments on the manuscript.

## Literature cited

Alekseyenko, A. A., A. A. Gorchakov, B. M. Zee, S. M. Fuchs, P. V. Kharchenko et al., 2014 Heterochromatin-associated interactions of Drosophila HP1a with dADD1, HIPP1, and repetitive RNAs. Genes Dev. 28: 1445–60.

Aruna, S., H. A. Flores, and D. A. Barbash, 2009 Reduced fertility of Drosophila melanogaster hybrid male rescue (Hmr) mutant females is partially complemented by Hmr orthologs from sibling species. Genetics. 181: 1437–50.

Barbash, D. A., 2010 Genetic testing of the hypothesis that hybrid male lethality results from a failure in dosage compensation. Genetics 184: 313–16.

Benna, C., S. Bonaccorsi, C. Wulbeck, C. Helfrich-Forster, M. Gatti et al., 2010 Drosophila timeless2 is required for chromosome stability and circadian photoreception. Curr. Biol. 20: 346– 352.

Bhat, M. A., A. V. Philp, D. M. Glover, and H. J. Bellen, 1996 Chromatid segregation at anaphase requires the barren product, a novel chromosome-associated protein that interacts with topoisomerase II. Cell 87:1103–14.

Bolkan B. J., R. Booker, M. L. Goldberg, and D. A. Barbash, 2007 Developmental and cell cycle progression defects in Drosophila hybrid males. Genetics. 2007 177: 2233–41.

Bonaccorsi, S., M. G. Giansanti, and M. Gatti, 2000 Spindle assembly in Drosophila neuroblasts and ganglion mother cells. Nat. Cell Biol. 2: 54–56.

Cenci, G., R. B. Rawson, G. Belloni, D. H. Castrillon DH, M. Tudor et al., 1997 UbcD1, a Drosophila ubiquitin-conjugating enzyme required for proper telomere behavior. Genes Dev. 11: 863–75.

Cicconi, A., E. Micheli, F. Vernì, A. Jackson, A. C. Gradilla, et al., 2017 The Drosophila telomere-capping protein Verrocchio binds single-stranded DNA and protects telomeres from DNA damage response. Nucleic Acids Res. 45:3068–85

Chang, H-J., S. Goulding, Earnshaw W. C., and M. Carmena, 2003 RNAi analysis reveals an unexpected role for topoisomerase II in chromosome arm congression to a metaphase plate. J Cell Sci. 116: 4715–26.

Cobbe, N., E. Savvidou and M. M. Heck, 2006 Diverse mitotic and interphase functions of condensins in Drosophila. Genetics 172:991–1008.

Coelho, P. A., J. Queiroz-Machado J, A. M. Carmo, S. Moutinho-Pereira S, H. Maiato et al., 2008 Dual role of topoisomerase II in centromere resolution and aurora B activity. PLoS Biol. 6: e207.

Coelho, P. A., J. Queiroz-Machado, and C. E. Sunkel, 2003 Condensin-dependent localisation of topoisomerase II to an axial chromosomal structure is required for sister chromatid resolution during mitosis. J Cell Sci. 116: 4763–76.

Dorsett, D., and L. Ström, 2012 The ancient and evolving roles of cohesin in gene expression and DNA repair. Curr Biol. 22:R240–50.

Durante, M., J. S. Bedfordì, D. J. Chen, S. Conrad, M. N. Cornforth, et al., 2013 From DNA damage to chromosome aberrations: joining the break.Mutat Res. 756:5–13.

Elgin, S. C., and G. Reuter, 2013 Position-effect variegation, heterochromatin formation, and gene silencing in Drosophila. Cold Spring Harb Perspect Biol. 5: a017780.

Fanti, L., G. Giovinazzo, M. Berloco, and S. Pimpinelli, 1998 The heterochromatin protein 1 prevents telomere fusions in Drosophila. Mol Cell. 2: 527–38.

Fluminhan, A., and T. Kameya, 1996 Behaviour of chromosomes in anaphase cells in embryogenic callus cultures of maize (Zea mays L.) Theor. Appl. Genet. 92: 982–90

Gatti, M., 1979 Genetic control of chromosome breakage and rejoining in Drosophila melanogaster: spontaneous chromosome aberrations in X-linked mutants defective in DNA metabolism. Proc. Natl. Acad. Sci. USA 76: 1377–1381.

Gatti, M., and B. S. Baker, 1989 Genes controlling essential cell cycle functions in Drosophila melanogaster. Genes Dev. 3: 438–453.

Gatti, M., and M. L. Goldberg, 1991 Mutations affecting cell division in Drosophila. Methods Cell Biol. 35: 543–586.

Gatti, M. and S. Pimpinelli, 1992 Functional elements in Drosophila melanogaster heterochromatin. Annu Rev Genet. 26:239–75.

Gatti, M., C. Tanzarella, and G. Olivieri, 1974 Analysis of the chromosome aberrations induced by x-rays in somatic cells of Drosophila melanogaster. Genetics 77: 701–719.

Greil, F., E. de Wit, H.J. Bussemaker and B. van Steensel, 2007, HP1 controls genomic targeting of four novel heterochromatin proteins in Drosophila. EMBO J. 26: 741–51.

Joppich, C., S. Scholz, G. Korge, and A. Schwendemann, 2009 Umbrea, a chromo shadow domain protein in Drosophila melanogaster heterochromatin, interacts with Hip, HP1and HOAP Chromosome Res. 17:19–36

Heeger, S., O. Leismann, R. Schittenhelm, O. Schraidt, S. Heidmann, et al., 2005 Genetic interactions of separase regulatory subunits reveal the diverged Drosophila Cenp-C homolog. Genes Dev. 19: 2041–53

Hirano, T., 2015 Chromosome Dynamics during Mitosis. Cold Spring Harb Perspect Biol. 7: a015792

Hoskins, R. A., C. D. Smith, J. W. Carlson, A. B. Carvalho, A. Halpern, et al., 2002 Heterochromatic sequences in a Drosophila whole-genome shotgun assembly. Genome Biol. 3: RESEARCH0085

Lattao, R., S. Bonaccorsi, X. Guan, S. A. Wasserman, and M. Gatti, 2011 Tubby-tagged balancers for the Drosophila X and second chromosomes. Fly (Austin) 5: 369–370.

Maheshwari, S, and D. A. Barbash, 2011, The genetics of hybrid incompatibilities. Annu Rev Genet. 45: 331–55.

Maheshwari, S., and D. A., Barbash, 2012 Cis-by-Trans regulatory divergence causes the asymmetric lethal effects of an ancestral hybrid incompatibility gene. PLoS Genet. 8: e1002597

Marzio, A., C. Merigliano, M. Gatti, and F. Vernì, 2014 Sugar and chromosome stability: clastogenic effects of sugars in vitamin B6-deficient cells. PLoS Genet. 10: e1004199.

Mengoli, V., E. Bucciarelli, R. Lattao, R. Piergentili, M. Gatti, et al., 2014 The analysis of mutant alleles of different strength reveals multiple functions of topoisomerase 2 in regulation of Drosophila chromosome structure. PLoS Genet. 10: e1004739.

Merigliano, C., A. Marzio, F. Renda, M. P. Somma, M. Gatti, et al., 2017 A Role for the Twins Protein Phosphatase (PP2A-B55) in the Maintenance of Drosophila Genome Integrity. Genetics 205:1151–67.

Miklos G. L., and J. N. Cotsell, 1990 Chromosome structure at interfaces between major chromatin types: alpha- and beta-heterochromatin. Bioessays 12:1–6.

Obe, G., P. Pfeiffer, J. R. Savage, C. Johannes, W. Goedecke et al., 2002 Chromosomal aberrations: formation, identification and distribution. Mutat Res. 504:17–36

Oliveira, R. A., S. Kotadia, A. Tavares, M. Mirkovic, K. Bowlin, et al., 2014 Centromere-independent accumulation of cohesin at ectopic heterochromatin sites induces chromosome stretching during anaphase. PLoS Biol. 12:e1001962.

Orr, H. A., L. D. Madden, J. A. Coyne, R. Goodwin and R. S. Hawley, 1997 The developmental genetics of hybrid inviability: a mitotic defect in Drosophila hybrids. Genetics 145: 1031–1040.

Presgraves, D. C., 2010 The molecular evolutionary basis of species formation. Nat Rev Genet. 11:175–80.

Raffa, G.D., G. Siriaco, S. Cugusi, L. Ciapponi, G. Cenci, et al., (2009) The Drosophila modigliani (moi) gene encodes a HOAP-interacting protein required for telomere protection. Proc. Natl. Acad. Sci. U.S.A. 106: 2271–2276.

Raffa,G. D., D. Raimondo, C. Sorino, S. Cugusi, G. Cenci, et al., 2010 Verrocchio, a Drosophila OB fold-containing protein, is a component of the terminin telomere-capping complex. Genes Dev. 24: 1596–1601.

Ross B. D., L. Rosin, A. W. Thomae, M. A Hiatt, D. Vermaak, et al., 2013 Stepwise evolution of essential centromere function in a Drosophila neogene. Science. 340: 1211–4.

Satyaki, P.R., T. N. Cuykendall, K. H. Wei, N. J. Brideau, H. Kwak et al., 2014 The Hmr and Lhr hybrid incompatibility genes suppress a broad range of heterochromatic repeats. PLoS Genet. 10: e1004240.

Savvidou, E., N. Cobbe, S. Steffensen, S. Cotterill and M. M. Heck, 2006 Drosophila CAP-D2 is required for condensin complex stability and resolution of sister chromatids. J Cell Sci. 2005 Jun 1;118(Pt 11):2529–43.

Shimizu, N., K. Shingaki, Y. Kaneko-Sasaguri, T. Hashizume, and T. Kanda, 2005 When, where and how the bridge breaks: anaphase bridge breakage plays a crucial role in gene amplification and HSR generation Exp. Cell Res. 302: 233–43

Smith, D. A., B. S. Baker and M. Gatti, 1985 Mutations in genes encoding essential mitotic functions in Drosophila melanogaster. Genetics 110: 647–70

Somma, M. P., F. Ceprani, E. Bucciarelli, V. Naim, V. De Arcangelis et al., 2008 Identification of Drosophila mitotic genes by combining co-expression analysis and RNA interference. PLoS Genet. 4: e1000126.

Somma, M. P., B. Fasulo, G. Siriaco, and G. Cenci, 2003 Chromosome condensation defects in barren RNA-interfered Drosophila cells. Genetics. 165:1607–11.

Steffensen, S., P. A. Coelho, N. Cobbe, S. Vass, M. Costa, et al., 2001 A role for Drosophila SMC4 in the resolution of sister chromatids in mitosis. Curr Biol. 11:295–307

Thomae, A. W., G. O. Schade, J. Padeken, M. Borath, I. Vetter I, et al., 2013 A pair of centromeric proteins mediates reproductive isolation in Drosophila species. Dev Cell. 27: 412–24.

Uhlmann, F., 2016 SMC complexes: from DNA to chromosomes. Nat Rev Mol Cell Biol. 7:399–412.

Vermaak, D., and H. S. Malik, 2009 Multiple roles for heterochromatin protein 1 genes in Drosophila. Annu Rev Genet 43: 467–92.

Wei, Y., L. Yu, J. Bowen, M. A. Gorovsky, and C. D. Allis, 1999 Phosphorylation of histone H3 is required for proper chromosome condensation and segregation. Cell 97: 99–109

Williams, B. C., T. L. Karr, J. M. Montgomery, and M. L. Goldberg, 1992 The Drosophila l(1)zw10 gene product, required for accurate mitotic chromosome segregation, is redistributed at anaphase onset. J. Cell Biol. 118: 759–73.

Williams, B., G. Leung, H. Maiato A. Wong, Z. Li Z, et al., 2007 Mitch a rapidly evolving component of the Ndc80 kinetochore complex required for correct chromosome segregation in Drosophila. J Cell Sci. 120: 3522–33

Zhang, Y., L. Zhang, X. Tang, S. R. Bhardwaj, J. Ji, et al., 2016 MTV, an ssDNA Protecting Complex Essential for Transposon-Based Telomere Maintenance in Drosophila. PLoS Genet. 12: e1006435.

